# Ophiostomatalean fungi associated with *Cyrtogenius luteus* infested *Pinus massoniana*, including characterization of *Masuyamyces qingyuanensis* sp. nov. in China

**DOI:** 10.1101/2024.11.06.622303

**Authors:** Kun Liu, Xiang Gao, Ya-Jie Wang, Ming-Liang Yin

## Abstract

Ophiostomatalean fungi refers to the species in the order Ophiostomatales of Ascomycota. This group of fungi comprises wood blue-stain agents and some species that are important pathogens of trees, nematodes, or humans. Most species in the Ophiostomatales have beneficial relationships with wood-boring insects such as bark beetles. Recent surveys were conducted to investigate the diversity of Ophiostomatoid fungi associated with *Cyrtogenius luteus* infesting *Pinus massoniana* in some areas of Guangdong, China. This study, 382 fungal strains were obtained, and 224 were identified as belonging to the Ophiostomatales order. Identification of these strains revealed that they spanned seven distinct genera. A new species, *Masuyamyces qingyuanensis sp. nov*., was discovered and identified through a combination of morphological and phylogenetic analyses across four gene regions: beta-tubulin (TUBB), internal transcribed spacer (ITS), large subunit ribosomal RNA (LSU), and translation elongation factor 1 alpha (EF1A).

## Introduction

Ophiostomatoid fungi refers to *Ophiostoma* and *Ceratocystis* species that commonly co-occur in ecological niches associated with insects (Wingfield *et al*. 1993). The growth of some of these fungal associates can cause sapwood discoloration, leading to reduced profitability of the product; Ophiostomatoid fungi, due to their transmission mechanisms, are likely to increase in number in the coming years, increasing the likelihood that they will become one of the most critical threats to the world’s agricultural, forestry, and natural forest ecosystems (Wingfield *et al*. 2013). In the past, the classification of Ophiostomatales has been confused by the convergence of morphological characters. Fortunately, advances in molecular biology techniques such as gene sequencing and phylogeny have significantly changed the taxonomic classification of Ophiostomatales species. De Beer *et al*. (2022) reevaluated the generic boundaries of Ophiostomatales based on four gene regions (LSU, ITS, EF1A, and RPBII) from over 200 ex-type isolates in Ophiostomatales.

*Cyrtogenius luteus*, native to Asia, typically targets dying or dead trees (Gómez *et al*. 2017). This species has been documented in various provinces across China, including Guangdong and Fujian, and has a broad host range encompassing conifers such as *P. massoniana* and *P. tabuliformis* (Reference). Bark beetles parasitize conifers and always carry a variety of fungi, including Ophiostomatoid fungi (Kirisits 2004). Bark beetles form specialized fungal structures called “mycangia” (Batra 1963) that not only serve a transport function but also provide a medium for the fungus (Happ 1971). Ophiostomatalean fungi can produce sticky droplets on elevated ascospore necks or conidia, which makes them particularly suitable for arthropod transmission (De Beer *et al*. 2013). However, the diversity of ophiostomatoid fungi on *Pinus massoniana* is limited in Guangdong. This study aimed to explore the diversity of Ophiostomatales fungi on *P. massoniana* in Qingyuan and Zhaoqing, Guangdong Province. The research is anticipated to be instrumental in advancing our understanding of these fungi’s biodiversity and geographical distribution.

## Materials and Methods

### Sample Collection and Fungi Isolation

Between October 2021 and October 2022, we collected bark and blue-stained sapwood of *P. massoniana* affected by pine wood nematode disease in Zhaoqing (23°05’N, 112°44’E) and Qingyuan (23°7’N, 113°01’E), located in northwestern Guangdong Province. The bark beetles were collected in sterile centrifuge tubes, while the bark and wood samples were placed into sterile sampling bags. These samples were then stored at a temperature of 4°C. Within one week, we isolated fungi from these samples using somatic microscopy. Fungi isolated from the samples were inoculated onto malt extract agar (MEA: 20 g Biolab malt extract, 20 g Biolab agar, 1000 mL deionized water) plates, each with a diameter of 90 mm. Following inoculation, the plates were incubated at 25°C for one week in darkness. Subsequently, the mycelial tips from the resulting fungal colonies were sub-cultured onto 2% selective malt extract agar (MEAS: 0.5 g injectable Streptomycin Sulfate per liter of MEA).

The model cultures of Ophiostomatales fungi described in this study were maintained at the Chinese General Microbial Strain Collection and Management Center (CGMCC; http://www.cgmcc.net/english/catalogue.html), Beijing, China. The names of the new taxa were registered in MycoBank (http://www.mycobank.org/)

### DNA Extraction, PCR, and Sequencing

DNA extraction was followed methods by Yin et al. (2016). The primers were ITS5/ITS4 (White *et al*. 1990) for ITS; LROR/LR5 for LSU (Vilgalys & Hester 1990), EF2F (Marincowitz *et al*. 2015)/EF2R (Jacobs *et al*. 2004) for EF1A, and bt2a/bt2b (Glass & Donaldson 1995) for TUBB.

Each PCR reaction consisted of 25 μL, comprised of 12.5 μL of 2×Taq Master Mix (buffer, dNTPs, and Taq; Thermo Fisher Scientific), 0.5 μL of both forward and reverse primers, 10.5 μL of PCR-grade deionized water, and 1 μL of DNA template. The PCR amplification procedure followed these steps: initial denaturation at 95°C for 3 minutes; then 35 cycles of denaturation at 95°C for 30 seconds, annealing at 52-55°C for 30 seconds, and extension at 72°C for 1 minute; a final extension at 72°C for 10 minutes; and a final hold at 12°C. All PCR products were sequenced in Sangong Biotechnology Co., Guangzhou, China. Sequences were assembled using Geneious v. 7.1.4 (Biomatters, Auckland, New Zealand). Sequence data were identified using the BLAST algorithm from NCBI GenBank (Altschul *et al*. 1990). Sequences used for phylogenetic analyses were submitted to GenBank (Table 1).

**Table 1.**
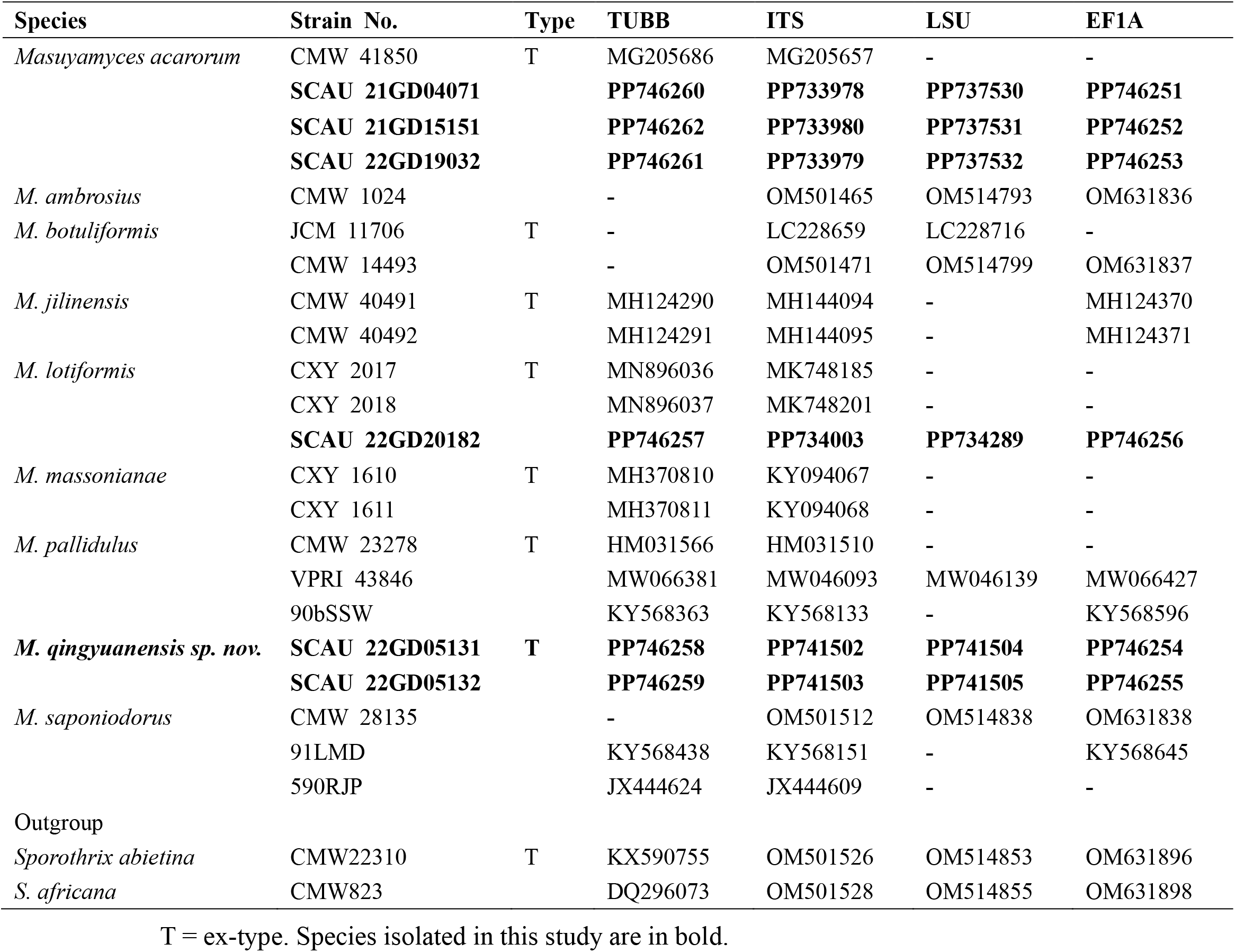
Isolates were used for phylogenetic analyses in this study.

### Phylogenetic Analyses

Initial strain identification was conducted using standard BLAST searches. Due to the wide variation in gene regions for Ophiostomatales fungi and species complexes across public databases, we referred to previous studies to select the appropriate gene regions required for constructing the phylogenetic tree. For phylogenetic analysis, distinct datasets were compiled for each gene region: ITS, LSU, TUBB, and EF1A. These datasets included sequences generated in our study and sequences obtained from BLAST searches of acquired strains in the NCBI GenBank database. Additionally, representative sequences with the highest similarity scores and model strain sequences of closely related species were downloaded for analysis. The corresponding phylogenetic trees display the accession numbers for these sequences (Fig. 1). Each dataset was compiled using MEGA X software (Kumar *et al*. 2018). We utilized the online tool MAFFT v.7 for sequence alignment, which is accessible at https://mafft.cbrc.jp/alignment/server/ (Katoh & Standley 2013)—and using the AUTO (FFT-NS-1, FFT-NS-2, FFT-NS-i or L-INS-i; depends on data size) strategy for the ITS dataset and the E-INS-i strategy and default settings for the other datasets.

**FIGURE 1.**
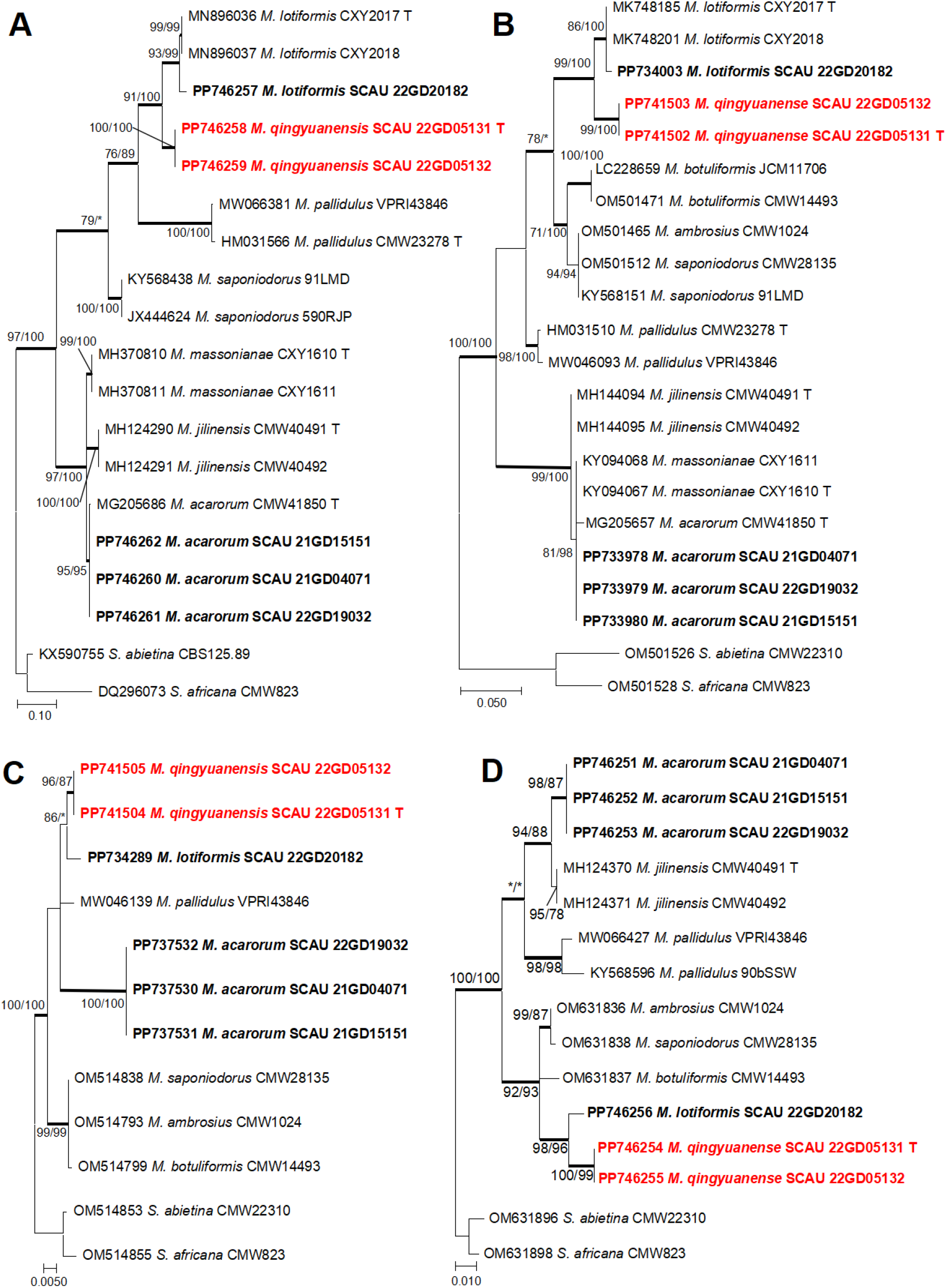
ML tree of *Masuyamyces*. A: generated from the *β-tub* sequence data. B: generated from the ITS sequence data. C: generated from the LSU sequence data. D: generated from the *TEF1-α* sequence data. Sequences generated from this study are printed in bold. Bold branches indicate posterior probability values ≥ 0.9. Bootstrap values ≥ 70 % are recorded as ML/MP at the nodes. * = Bootstrap values < 70 %.

**FIGURE 2.**
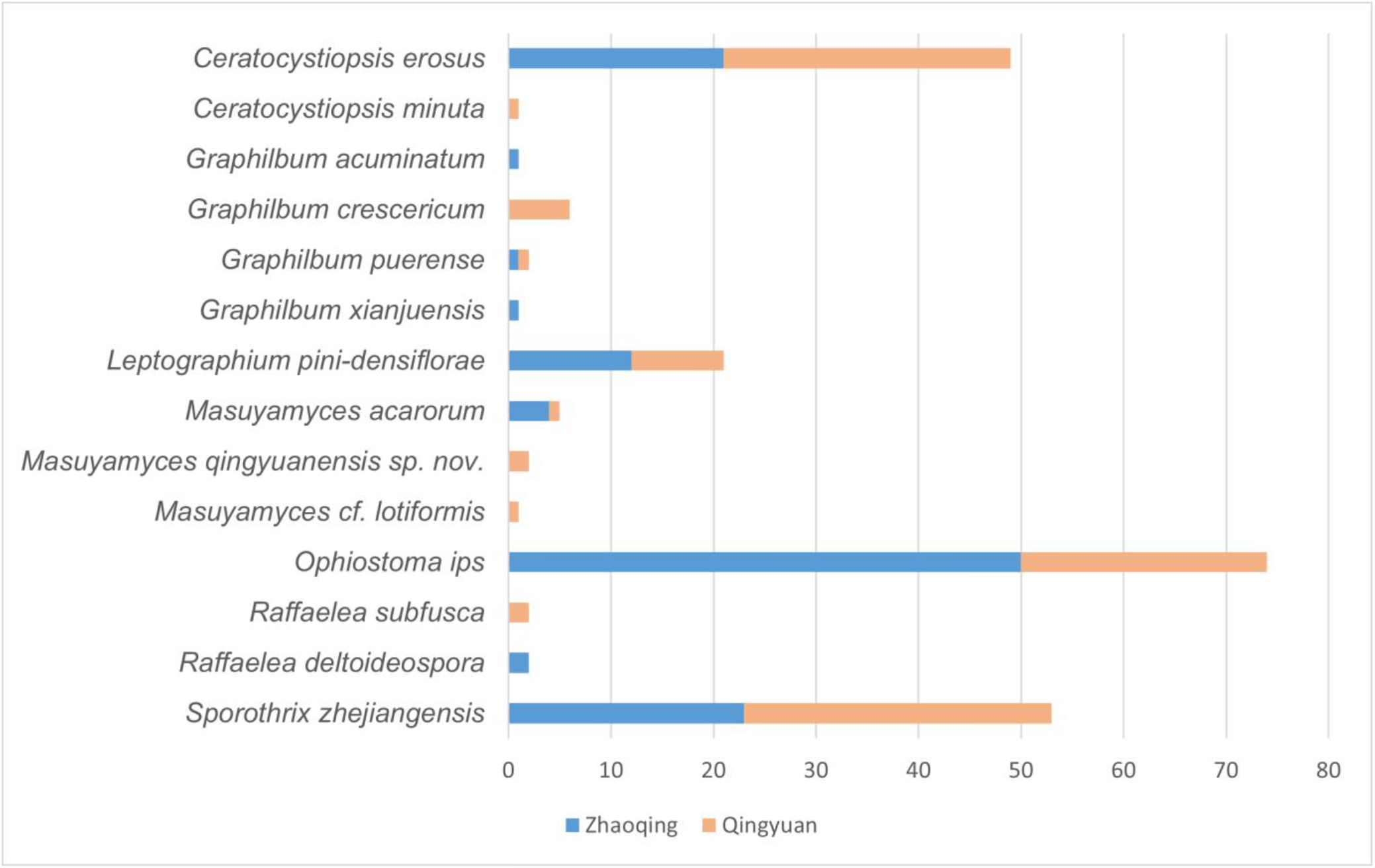
The distribution of isolated fungi of Ophiostomatales in this study.

The best alternative model for each dataset was found using Jmodeltest v. 2.1.10 (Darriba *et al*. 2012), the HKY+G model was selected for the *β-tub* datasets, the GTR+G model for the ITS datasets, and the GTR+I model for the LSU and *TEF1-α* datasets. Maximum likelihood (ML) analyses were performed separately for the different datasets using MEGA-X with 1000 bootstrap replicates. Maximum parsimony (MP) analysis was performed with PAUP* version 4.0b10 (Swofford 2002). Bayesian inference (BI) was carried out with MrBayes-3.2.7-WIN (Ronquist *et al*. 2012), employing four Markov Chain Monte Carlo (MCMC) chains that ran simultaneously from a tree initialized at random. The chains were run for five million generations, with a sample taken every 100 generations. During the ‘burn-in’ phase, 25% of sampled trees were discarded to account for the startup bias. Posterior probabilities were then computed from the remaining trees. The results were visualized and analyzed using FigTree v1.4.4. Table 2 summarizes the statistical values of the relevant parameters for all datasets utilized in the phylogenetic analysis.

**TABLE 2.** Parameters used and statistical values related to all phylogenetic analyses in the present study.

|  |  | TUBB | ITS | LSU | EF1A |
| --- | --- | --- | --- | --- | --- |
| Alignments | Number of taxa | 19 | 22 | 12 | 15 |
|  | Total | 588 | 526 | 794 | 416 |
|  | Constant | 97 | 295 | 741 | 357 |
|  | Uninformative | 94 | 53 | 10 | 10 |
|  | Informative | 397 | 178 | 43 | 49 |
| MP | Number of trees | 1 | 1 | 4 | 3 |
|  | Tree length | 870 | 370 | 53 | 72 |
|  | CI | 0.787 | 0.757 | 0.849 | 0.750 |
|  | RI | 0.919 | 0.894 | 0.920 | 0.867 |
|  | RC | 0.724 | 0.677 | 0.781 | 0.650 |
|  | HI | 0.213 | 0.243 | 0.151 | 0.250 |
| Model tests | Subst. model | HKY+G | GTR+G | GTR+I | GTR+I |
| ML | P-inv | - | - | 0.470 | 0.430 |
|  | Gamma | 0.408 | 0.293 | - | - |
| BI | Burn-in | 25% | 25% | 25% | 25% |

### Morphological studies

A model culture and another isolate were selected for culture studies. An agar block of approximately 5 mm diameter was removed from the edge of an actively growing fungal colony and placed in the center of a 90 mm diameter MEA to study growth rate, incubated at 25°C in the dark. Colony diameters measured by crisscrossing were recorded on days 7 and 14 until the fastest-growing mycelium reached the edge of the MEA.

For the microstructural study of the new species, the strains were inoculated on pine strip plates (20 g of agar powder in 1000 ml of deionized water, to which two sterilized pine strips were added just before solidification) and incubated at 25°C in the dark with daily observations. After 14 days of incubation on pine strip plates, slides were made to observe sexual/asexual structures. The microstructures of Ophiostomatoid fungi were measured and photographed using a Zeiss Axioscope 5 (Carl Zeiss, Germany). Thirty measurements were taken for each taxonomically informative structure, such as conidiophores and conidial peduncles, and measurements were expressed in the format of (min-) mean minus standard deviation - mean plus standard deviation (-max).

## Results

### Fungal isolation

In this study, 19 samples were collected from Zhaoqing, yielding 187 fungal isolates, of which 119 were identified as Ophiostomatales fungi. Additionally, 23 samples were collected from Qingyuan, resulting in 195 fungal isolates, including 105 Ophiostomatales fungi. Preliminary identification of the Ophiostomatales fungi isolated in this study revealed the following distribution: *Ceratocystiopsis* (50 isolates), *Graphilbum* (11 isolates), *Leptographium* (21 isolates), *Masuyamyces* (9 isolates), *Ophiostoma* (73 isolates), *Raffaelea* (2 isolates), *Sporothrix* (53 isolates), and five isolates that remained unidentified.

### Phylogenetic Analyses

Based on the sequence data of Masuyamyces obtained from the four gene regions, phylogenetic analysis revealed that *M. qingyuanensis* consistently formed well-supported, independent branches across all four phylogenetic trees.

### Taxonomy

***Masuyamyces qingyuanensis*** K. Liu & M.L. Yin, *sp. nov*.

(Fig. 3)

**FIGURE 3.**
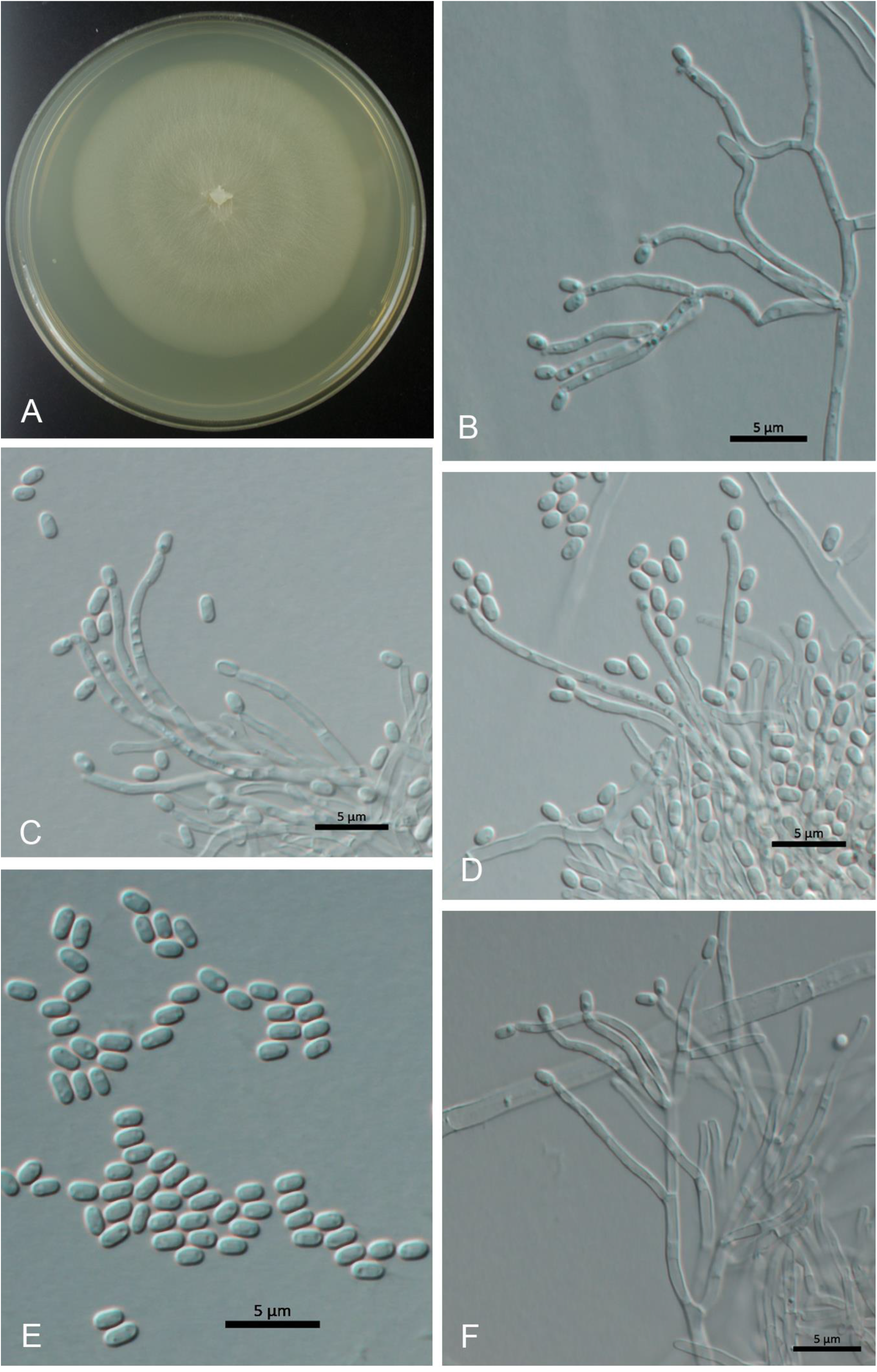
*Masuyamyces qingyuanensis sp. nov*. A: Pure culture gown on MEA in the dark for 14 days; B – D, F: Conidiogenous cells giving rise to oblong conidia; E: Conidia; scale bars: 5 μm.

MycoBank 853760

**Etymology**. The name is derived from the toponymic name of the collection site.

**Type**. Dinghu mountain, Qingyuan, Guangdong Province, China, Oct. 2022, from the galleries of *P. massoniana*, 300m elevation, holotype HAMS 352761, ex-holotype culture SCAU 22GD05131 = CGMCC 3.27006.

Sexual state not observed. Asexual stage: sporangia forming on nutrient mycelium, conidiophore hyalorhinocladiella-like, clustered, conidia arising singly or in pairs from the apex, cylindrical, hyaline, septate, measuring (1.4-) 1.5 - 1.7 (−1.9) × 0.8 -1.0 μm.

Cultural characteristics. Colonies on MEA initially appeared as a thin white layer.

The mycelium spread outward, with the edges not being flat. Under dark conditions at a constant temperature of 25°C, the fungal growth can reach up to 69 mm in diameter within 14 days. During this period, distinct ring marks appear on the colony, and the edges of the colony become smooth.

**Host tree:** *Pinus massoniana*.

**Distribution:** China

**Notes:** This species is closely related to *M. lotiformis*; however, its conidia are significantly smaller. While the sexual morphology of *M. lotiformis* remains unknown, the asexual morphology is Hyalorhinocladiella-like, characterized by hyphal cells produced from surface hyphae. The conidia are hyaline, indehiscent, smooth, and have a clavate to ovate shape, with dimensions ranging from (3.5 -)4-5.5(−6.5) × (2 -)2.5-3.5(−4) μm. On 2% malt extract agar (MEA), colonies grew to a diameter of 65 mm within 15 days at 25°C, exhibiting white coloration and smooth edges. The optimal temperature for growth was determined to be 30°C (Wang *et al*. 2020). *M. qingyuanensis* and *M. lotiformis* have high support for forming separate branches on all four phylogenetic trees.

## Discussion

This study discovered a relatively high abundance of Ophiostomatales fungi species in Guangdong. The genera *Ophiostoma, Sporothrix*, and *Ceratocystiopsis* were particularly prevalent, representing 46.07% of all strains isolated in this research. *O. ips* stood out as the most frequently isolated strain. The findings from this study’s fungal isolation efforts have expanded our understanding of the diversity of Ophiostomatalean fungi in Guangdong Province. Furthermore, based on morphological and phylogenetic analyses, *M. qingyuanensis* was identified and recognized as a new species. Yin *et al*. (2019) suggest that integrating multiple lines of evidence, such as morphology, DNA, substratum, and geography, represents the progressive approach to fungal taxonomy.

## Acknowledgment

This study was funded by the National Natural Science Foundation of China (32070012) and the Guangdong Basic and Applied Basic Research Foundation (2020A1515010486, 2022A1515010901). We would like to thank all the forestry staff who assisted us during the sampling process in Guangdong Provinces over the years; we especially appreciate Professor J. Wang for his unwavering support and care for our laboratory-related work.

## Declarations

### Availability of data and material

The manuscript mentioned all the availability of data. Ex-type cultures of new species were deposited in the Culture Collection of South China Agricultural University (SCAU) and the China General Microbiological Culture Collection Center (CGMCC). The type herbariums were preserved in the Fungarium (HAMS), Institute of Microbiology, Chinese Academy of Sciences. DNA sequence data are available in Genebank (https://www.ncbi.nlm.nih.gov/nucleotide/), and taxonomic novelties are available in Mycobank (https://www.mycobank.org).

### Authors’ contributions

Conceptualization, Mingliang Yin; Data curation, Kun Liu; Funding acquisition, Mingliang Yin; Formal analysis, Kun Liu; Funding acquisition, Mingliang Yin; Investigation, Xiang Gao, Kun Liu, and Yajie Wang; Methodology, Mingliang Yin; Project administration, Mingliang Yin; Resources, Xiang Gao, Kun Liu, Minjie Chen, and Yajie Wang; Software, Kun Liu; Supervision, Mingliang Yin; Validation, Mingliang Yin; Visualization, Mingliang Yin; Writing – original draft, Kun Liu; Writing – review & editing, Mingliang Yin.

## References

Altschul, S.F., Gish, W., Miller, W., Myers, E.W. & Lipman, D.J. (1990) Basic local alignment search tool. Journal of Molecular Biology 215 (3): 403–410. 10.1016/S0022-2836(05)80360-2

Batra, L.R. (1963) Ecology of Ambrosia fungi and their dissemination by beetles. Transactions of the Kansas Academy of Science 66 (2): 213–236. 10.2307/3626562

Darriba, D., Taboada, G.L., Doallo, R. & Posada, D. (2012) jModelTest 2: more models, new heuristics and parallel computing. Nature Methods 9 (8): 772. 10.1038/nmeth.2109

De Beer, Z.W., Seifert, K.A. & Wingfield, M.J. (2013) The ophiostomatoid fungi: their dual position in the Sordariomycetes. In: Seifert, K.A., De Beer, Z.W. & Wingfield, M.J. (Eds.) The Ophiostomatoid Fungi: Expanding Frontiers. CBS-KNAW Fungal Biodiversity Centre, Utrecht, The Netherlands, pp. 1–19.

De Beer, Z.W., Procter, M., Wingfield, M.J., Marincowitz, S. & Duong, T.A. (2022) Generic boundaries in the Ophiostomatales reconsidered and revised. Studies in Mycology 101: 57–120. 10.3114/sim.2022.101.02

Glass, N.L. & Donaldson, G.C. (1995) Development of primer sets designed for use with the PCR to amplify conserved genes from filamentous ascomycetes. Applied and Environmental Microbiology 61 (4): 1323–1330. 10.1128/aem.61.4.1323-1330.1995

Gómez, D.F., Hirigoyen, A., Balmelli, G., Viera, C.L. & Martínez, G. (2017) Patterns in flight phenologies of bark beetles (Coleoptera: Scolytinae) in commercial pine tree plantations in Uruguay. Bosque (valdivia) 38: 47–53. 10.4067/S0717-92002017000100006

Happ, G.M., Happ, C.M. & Barras, S.J. (1971) Fine structure of the prothoracic mycangium, a chamber for the culture of symbiotic fungi, in the southern pine beetle, Dendroctonus frontalis. Tissue and Cell 3 (2): 295–308. 10.1016/S0040-8166(71)80024-1

Jacobs, K., Bergdahl, D.R., Wingfield, M.J., Halik, S., Seifert, K.A., Bright, D.E. & Wingfield, B.D. (2004) Leptographium wingfieldii introduced into North America and found associated with exotic Tomicus piniperda and native bark beetles. Mycological Research 108 (4): 411–418. 10.1017/S0953756204009748

Katoh, K. & Standley, D.M. (2013) MAFFT multiple sequence alignment software version 7: improvements in performance and usability. Molecular Biology and Evolution 30 (4): 772–780. 10.1093/molbev/mst010

Kirisits, T. (2004) Fungal Associates of European Bark Beetles With Special Emphasis on the Ophiostomatoid Fungi. In: Lieutier, F., Day, K.R., Battisti, A., Grégoire, J.C. & Evans, H.F. (Eds.) Bark and Wood Boring Insects in Living Trees in Europe, a Synthesis. Kluwer Academic Publishers, Dordrecht, The Netherlands, pp.181–235. 10.1007/978-1-4020-2241-8_10

Kumar, S., Stecher, G., Li, M., Knyaz, C. & Tamura, K. (2018) MEGA X: Molecular Evolutionary Genetics Analysis across Computing Platforms. Molecular biology and evolution 35 (6): 1547–1549. 10.1093/molbev/msy096

Marincowitz, S., Duong, T.A., De Beer, Z.W. & Wingfield, M.J. (2015) Cornuvesica: A little known mycophilic genus with a unique biology and unexpected new species. Fungal Biology 119 (7): 615–630. 10.1016/j.funbio.2015.03.007

Ronquist, F., Teslenko, M., van der Mark, P., Ayres, D.L., Darling, A., Höhna, S., Larget, B., Liu, L., Suchard, M.A. & Huelsenback, J.P. (2012) MrBayes 3.2: efficient Bayesian phylogenetic inference and model choice across a large model space. Systematic Biology 61 (3): 539–542. 10.1093/sysbio/sys029

Swofford, D.L. (2002) PAUP*: Phylogenetic analysis using parsimony (*and other methods), Version 4.0b10.

Vilgalys, R. & Hester, M. (1990) Rapid genetic identification and mapping of enzymatically amplified ribosomal DNA from several Cryptococcus species. J Bacteriol 172 (8): 4238–4246. 10.1128/jb.172.8.4238-4246.1990

Wang, Z., Liu, Y., Wang, H.M., Meng, X.J., Liu, X.W., Decock, C., Zhang, X.Y. & Lu, Q. (2020) Ophiostomatoid fungi associated with Ips subelongatus, including eight new species from northeastern China. IMA Fungus 11: 3. 10.1186/s43008-019-0025-3

White, T.J., Bruns, T.D., Lee, S.B. & Taylor, J.W. (1990) Amplification and direct sequencing of fungal ribosomal RNA genes for phylogenetics. In: Innis, M.A., Gelfand, D.H., Sninsky, J.J. & White, T.J. (Eds.) PCR protocols: a guide to methods and application. Academic Press, San Diego, USA: 315–322. 10.1016/B978-0-12-372180-8.50042-1

Wingfield, M.J., Roux, J., Wingfield, B.D. & Slippers, B. (2013) Ceratocystis and Ophistoma: international spread, new associations and plant health. In: Seifert, K.A., De Beer, Z.W. & Wingfield, M.J. (Eds.) The Ophiostomatoid Fungi: Expanding Frontiers. CBS-KNAW Fungal Biodiversity Centre, Utrecht, The Netherlands, 12: 191– 200.

Wingfield, M.J., Seifert, K.A. & Webber, J.F. (1993) Ceratocystis and Ophiostoma: taxonomy, ecology and pathogenicity. American Phytopathological Society Press, St. Paul, MN, USA, 304 pp.

Yin, M.L., Wingfield, M.J., Zhou, X.D., Linnakoski, R. & De Beer, Z.W. (2019) Taxonomy and phylogeny of the Leptographium olivaceum complex (Ophiostomatales, Ascomycota), including descriptions of six new species from China and Europe. Mycokeys 60: 93–123. 10.3897/mycokeys.60.39069

